# Enhanced Single RNA Imaging Reveals Dynamic Gene Expression in Live Animals

**DOI:** 10.1101/2022.07.26.501631

**Authors:** Yucen Hu, Jingxiu Xu, Erqing Gao, Xueyuan Fan, Jieli Wei, Suhong Xu, Weirui Ma

**Author notes:** These authors contributed equally: Yucen Hu, Jingxiu Xu, and Erqing Gao. correspondence (Weirui Ma), (Suhong Xu).

## Abstract

Imaging endogenous mRNAs in live animals is technically challenging. Here we describe an MS2 based signal Amplification with Suntag System that enables live-cell RNA imaging of high temporal resolution and with 8xMS2 stem-loops, which overcomes the obstacle of inserting a 1,300 nt 24xMS2 into the genome for the imaging of endogenous mRNAs. Using this tool we were able to image the activation of gene expression and the dynamics of endogenous mRNAs in the epidermis of live *C. elegans*.

## Introduction

RNAs relay genetic information from DNA to proteins or function by themself. Live cell imaging of RNAs at a single molecule level is crucial to uncovering their roles in gene expression regulation^1^. Various tools have been developed to visualize RNAs in live cells^2, 3^, including RNA-binding protein – fluorescent protein approaches^4^, CRISPR-based systems^5, 6^, and those utilizing fluorophore-RNA aptamer pairs^7–9^. The MS2 based system is the most widely used and represents the current gold standard for single-molecule RNA imaging in live cells^2, 4^. MS2 is a short RNA stem-loop bound specifically by the bacteriophage MS2 coat protein (MCP). To image RNA, 24xMS2 are placed at the 3’UTR or 5’UTR, and a fluorescent protein is fused to the MCP (MCP-FP). When coexpressed in cells, up to 48 fluorescent proteins (2 X 24, with two MCPs bound to one MS2) will be recruited to the RNA through MS2-MCP binding. This forms a fluorescent spot indicating a single RNA molecule^2^.

The MS2 system has been successfully used to trace the whole mRNA life-cycle from transcription, to nuclear export, subcellular localization, translation, and to final degradation^2, 3^. However, most RNA imaging studies in animal cells have been performed using exogenous mRNAs in cultured cell lines. 24xMS2 have to be knocked into a specific genomic locus to image endogenous mRNA. The difficulty and low efficiency for the knock-in of long sequences into the genome represents a significant obstacle towards visualizing endogenous mRNA using the MS2 system. Thus, it is not surprising that less than ten endogenous mRNAs have been imaged in live animal cells at a single-molecule level, and examples of endogenous mRNAs imaged in live animals remain extremely rare^10–15^. Since overexpressed mRNAs may not faithfully recapitulate endogenous mRNA expression and dynamics. Development of more sensitive techniques for endogenous mRNA imaging is of great value.

## Results

In this study, we reasoned that combining the MS2 with a signal amplifier may allow the recruitment of more fluorescent proteins to the RNA with fewer MS2 repeats, (i.e. 8xMS2 - see Fig. 1a). To achieve this, we combined the MS2 and Suntag systems. Suntag is a 19 amino acid protein tag that binds to its specific single-chain variable fragment (scFv) antibody^16^. We fused MCP with a 24xSuntag array and linked scFv with sfGFP. When coexpressed in cells, one MS2 interacts with two MCP-24xSuntag molecules, further recruiting 2 × 24 sfGFP molecules (Fig. 1b). As the Suntag serves as a signal amplifier, the combined system was named as MS2 based signal Amplification with Suntag System (MASS). When an 8xMS2 is placed into the 3’UTR, up to 384 (2 × 8 × 24) sfGFP can then be tethered to a single mRNA through the MCP-Suntag-scFv-sfGFP interaction (Fig. 1a). This leads to the formation an intense GFP spot associated with single mRNA, facilitating live RNA imaging.

**Fig. 1.**
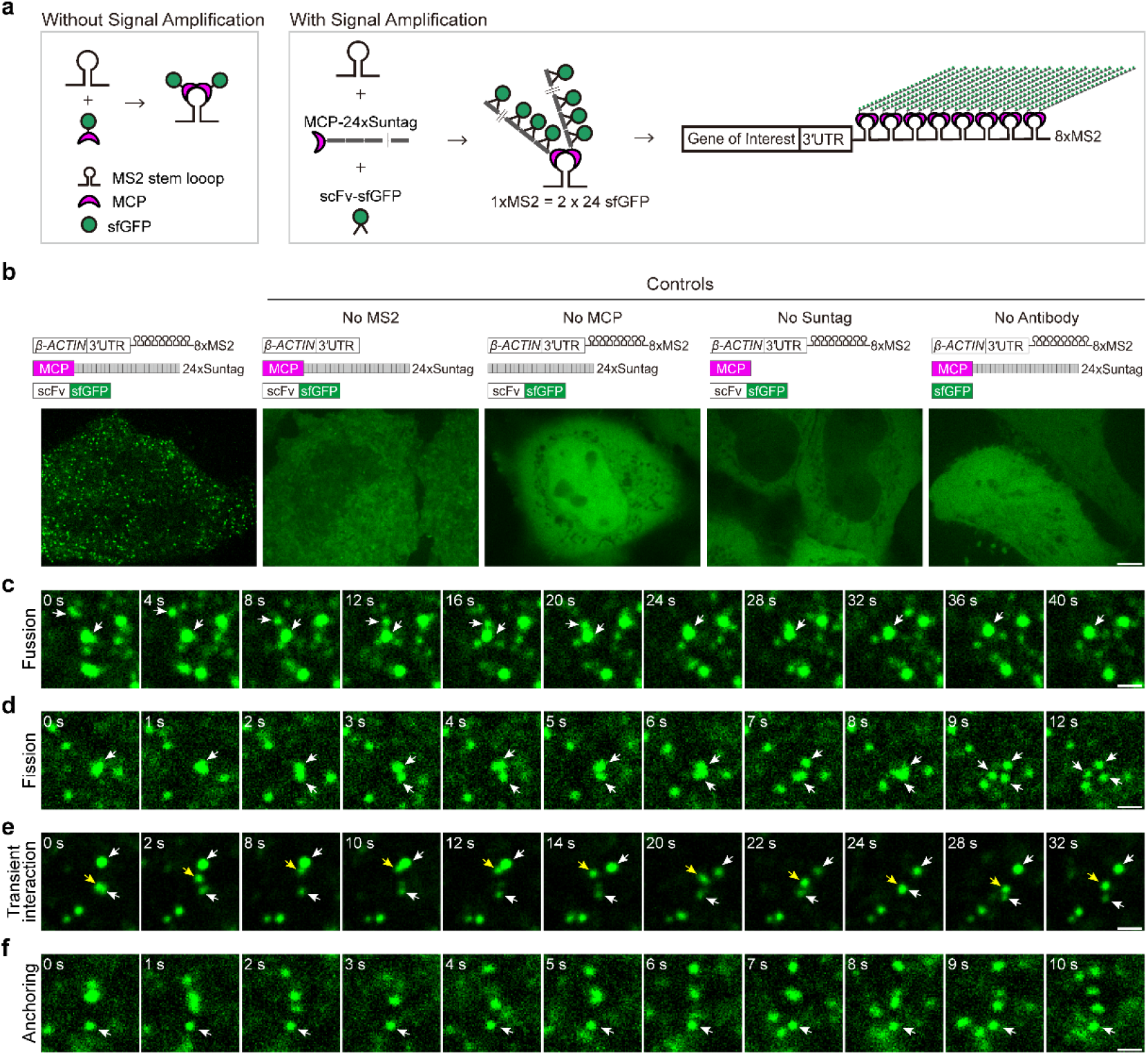
Live-cell imaging of *β-ACTIN* mRNA with the MS2 based signal amplification with Suntag System. **a**, Schematic of the classical MS2-MCP system and the MS2 based signal amplification with Suntag System. **b**, Representative images of *β-ACTIN-8xMS2* mRNA in live HeLa cells. The sfGFP fluorescence signal is shown. Left panel: Constructs of *β-ACTIN-8xMS2*, MCP-24xSuntag, and scFv-sfGFP were cotransfected into HeLa cells. Images were taken 12 hr after transfection. Right panels: where one of the elements was removed (as indicated). Scale bar, 5 μm. **c** to **f**, Time-lapse imaging of *β-ACTIN-8xMS2* mRNA dynamics in HeLa cells. sfGFP foci (*β-ACTIN* mRNAs) are shown. Constructs of *β-ACTIN-8xMS2*, MCP-24xSuntag, and scFv-sfGFP were cotransfected into HeLa cells. Images were taken 12 hr after transfection. **c**, A fusion event of two sfGFP spots (white arrows). **d**, A fission event: with large sfGFP foci split into three spots (white arrows). **e**, Transient interactions of an sfGFP spot (yellow arrow) between two spots (white arrows). **f**, An sfGFP spot showing no movement over a 10 second period. Scale bars: 1 μm.

As proof of concept, 8xMS2 was fused to the 3’UTR of *β-ACTIN* mRNA and transfected into HeLa cells. When all the required elements of the MASS (MS2, MCP, 24xSuntag, and scFv antibody) were present, bright GFP foci were readily detected (Fig. 1b). As controls, no GFP foci were detected when omitting any one of these elements (Fig. 1b). Thus, the GFP foci clearly represent *β-ACTIN* RNA molecules. It has also been reported that *β-ACTIN* mRNAs with 3’UTR can localize to the lamellipodia^17^. In support of this, we observed that the sfGFP foci of *β-ACTIN-3’UTR-8XMS2* mRNA were indeed localized to the lamellipodia in HeLa cells (Supplementary Fig. 1 and Supplementary Video 1). Taken together, these data indicated that MASS can be readlily used to image of RNA molecules in live cells without affecting RNA subcellular localization.

With 8xMS2 – 24xSuntag, a single mRNA molecule could be tethered with up to 384 sfGFP molecules, forming an sfGFP spot of high fluorescence intensity. Such an application therefore retains the ability to image mRNA using low power lasers, thus lowering any unwanted phototoxicity and photobleaching and still allowing the tracking of mRNA dynamics in high temporal resolution.

We then performed time-lapse imaging of the *β-ACTIN-3’UTR-8XMS2* mRNA with a time interval of 1 second in HeLa cells (Supplementary Video 2). We found that foci of mRNAs showed various dynamics; (1) Fusion. sfGFP spots fused into a more prominent spot (Fig 1c. and Supplementary Video 3); (2) Fission. Large sfGFP foci split into smaller spots (Fig. 1d and Supplementary Video 4); (3) Transient interaction. sfGFP foci touched each other briefly, then moved away (Fig. 1e and Supplementary Video 5), suggesting there are dynamic RNA-RNA interactions in cells; (4) Dynamic movement or anchoring. Despite most foci of *β-ACTIN-3’UTR-8XMS2* mRNAs showing dynamic movement in cells, other foci were far more static, showing little movement (Fig. 1f and Supplementary Video 6), suggesting these latter mRNAs may be anchored to subcellular structures.

Our ultimate goal was to develop tools for endogenous mRNA imaging in live animals. It has been previously reported that the knock-in of short sequences into the genome is far more efficient than those of longer sequences^18^. The MASS exploits this advantage as only 8xMS2 (350 nt) needs to be inserted into a genomic locus, thus overcoming the previous obstacle of the requirement of inserting a long 1,300 nt 24xMS2 into the genome for live-cell imaging of endogenous mRNA.

We then used the nematode *C. elegans* to specifically examine whether the MASS could be used for RNA imaging in live animals. An 8xMS2 was placed into the 3’UTR of *cdc-42* mRNA (Fig.2a). *cdc-42-8xMS2*, MCP-24xSuntag, and scFv-sfGFP were also expressed in the epidermis of live *C. elegans*. Consistent with the observations in HeLa cells, bright GFP foci could only be detected when all the required elements were present (Supplementary Fig. 2). Similalry, excluding any essential elements resulted in a complete failure of foci formation (Supplementary Fig. 2). Therefore, the MASS was efficient in imaging exogenous RNAs in live animals.

**Fig. 2.**
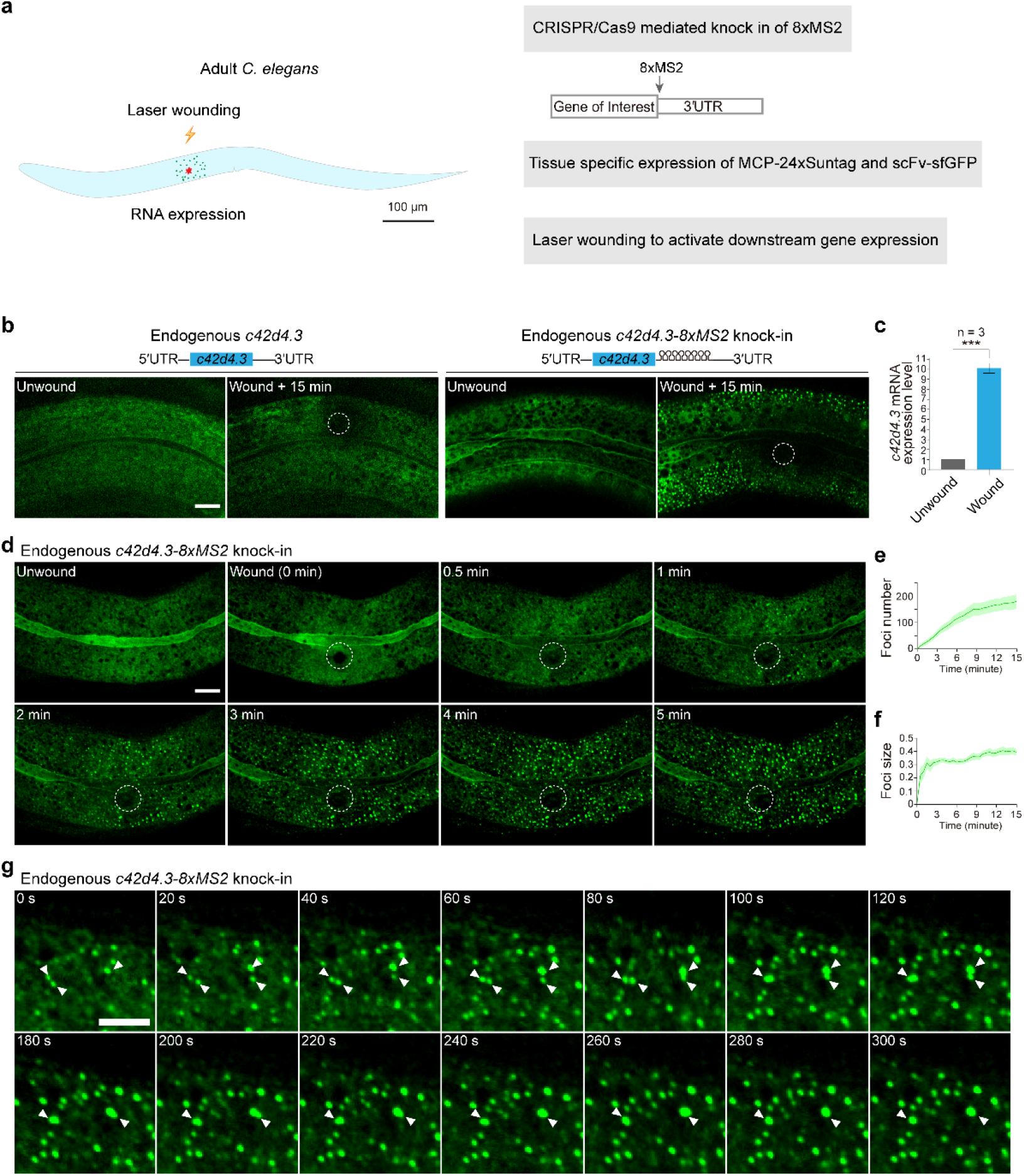
Live imaging of endogenous mRNAs in the epidermis of *C. elegans* using the MS2 based signal amplification with Suntag System. **a**, Schematic of the strategy for live imaging of endogenous mRNAs in the epidermis of *C. elegans*. **b**, Representative images of endogenous *c42d4.3-8xMS2* mRNA in the epidermis of live *C.elegans* using the strategy described in **a**. Left: *c42d4.3* without 8xMS2. Right: *c42d4.3* with 8xMS2. Images were taken before and 15 min after wounding. White dashed circles indicate wound sites. Scale bar, 10 μm. **c**, Quantitative RT-PCR showing the expression level of endogenous *c42d4.3* mRNA in *C. elegans* before and 15 min after wounding. *n* = 3 independent experiments; bars indicate mean ± SEM. Mann-Whitney test, ***p < 0.001. **d**, Time-lapse imaging of endogenous *c42d4.3-8xMS2* mRNA in the epidermis of live *C.elegans* before and after wounding. White dashed circles indicate the wound sites. Scale bar, 10 μm. **e** and **f**, shown are mean ± SEM of quantification of the number (**e**) and size (**f**) of sfGFP foci (endogenous *c42d4.3* mRNA) formed in the epidermis as measured 15 minutes after wounding. *n* = 10. **g**, Time-lapse imaging showing fusion of endogenous *c42d4.3* foci (white arrows) after laser wounding. Scale bar, 5 μm

Next, we set out to visualize gene expression activation and their dynamics of endogenous mRNAs in live animals. We used the skin of *C. elegans* as a model, which is composed of an epidermal epithelium with multiple nuclei. Upon wounding the epithelium via laser or needle, specific gene expressions and downstream signaling cascades for wound repair are then triggered and activated^19^. To this end, 8xMS2 was knocked into the 3’UTR region of two endogenous genes, *c42d4.3* and *mai-1* (Fig. 2b and Supplementary Fig. 3a). MCP-24xSuntag and scFv-sfGFP were expressed in the epidermis with the tissue-specific promoter *col-19*. We then used a UV laser to injure an area of the epidermis. Prior to wounding, no sfGFP foci were detected in either wild type or in 8xMS2 knock-in animals (Fig. 2b and Supplementary Fig. 3a). In constrct, numerous GFP foci were formed 15 min after wounding, in the *c42d4.3* and *mai-1* 8xMS2 knock-in animals but not in the wild type animals (Fig. 2b and Supplementary Fig. 3a). This suggests that the wounding had activated *c42d4.3* and *mai-1* mRNA expression. In agreement with this, qRT-PCR showed that *c42d4.3* and *mai-1* mRNA levels were upregulated more than eight-fold 15 min after injury (Fig. 2c and Supplementary Fig. 3b), confirming that GFP foci were able to detect endogenous *c42d4.3-8xMS2* and *mai-1-8xMS2* mRNAs.

Next, we tracked the dynamics of endogenous *c42d4.3-8xMS2* mRNA in the *C. elegans* epidermis after wounding. We found that in proximity to the injury site sfGFP foci were detected as early as 1 min after wounding (Fig. 2d, Supplementary Fig. 4, and Supplementary Video 7). As a control, we pretreated the *C. elegans* with Actinomycin D, which potently inhibits gene transcription. In such cases, no GFP foci could be detected after wounding (Supplementary Fig. 5). This indicated that sfGFP foci are newly synthesized *c42d4.3-8xMS2* mRNAs. Our data demonstrated that gene expression activation and transcription occur extremely fast (in this case, within 1 min after the stimulation). In addition, the formation of GFP foci gradually spreads from the injury site to distal regions. The total foci number steadily increased in the epidermis (Fig. 2d, 2e, Supplementary Fig. 4, and Supplementary Video 7). These data suggested that wounding generated a signal to activate downstream gene expression around the injury site. The signal was then diffused to distal regions and able to induce gene expression there. In addition, GFP foci preferentially underwent fusion leading to increased foci size (Fig. 2f, 2g, and Supplementary Video 8), suggesting that *c42d4.3* mRNAs undergo clustering after wounding and form RNA granules in vivo. In agreement with our observation, it has been previously reported that mRNAs formed large clusters and are co-translated in *Drosophila* embryos^12^.

## Discussion

It has been recently reported that a combination of PP7 and Suntag or Moontag facilitates long-term imaging of overexpressed mRNA in living cells^20^. However, whether it could be used to imaging endogenous RNA in live animals has not been tested. Here, we developed a tool to image RNA in live cells and animals using a signal amplification strategy. This utilized only 8xMS2 in comparison to the 24xMS2 used in the classic MS2-based live cell RNA imaging. The advantage of a short MS2 is of prime benifit for imaging endogenous mRNA as it overcomes the difficulties involved in inserting a long 1,300 nt 24xMS2 into a genomic locus. We expect this tool will help promote studies of RNA transcription, nuclear export, subcellular localization, translation, RNA sensing, and degradation, at the endogenous level in culture cells and live animals.

## Acknowledgments

We thank all members of the Ma lab for helpful discussions. We thank Christine Mayr (Memorial Sloan Kettering Cancer Center) for providing HeLa cells, pcDNA vectors, and Suntag constructs. We thank Jian Zhang (Yunnan University) for critically reading the manuscript and suggestions. This work was funded by the start-up funding from the Life Sciences Institute, the Leading innovation and entrepreneurship team of Hangzhou (TD2020006), the young fellows of Zhejiang University (2021QN81027) to W.M. and the National Key R&D Program of China (2021YFA1101002, 2021YFA1300302), the National Natural Science Foundation of China (91754111), and the Zhejiang Province Natural Science Foundation (2-2060203-21-001) to S.X.

## Author Contributions

W.M. and S.X. conceived the projects and designed the experiments. Y.H. performed all experiments and analyses in HeLa cells. J.X. and E.G. performed all experiments and analyses in *C. elegans*. X.F. provided help with cloning. J.W. provided help with cell culture. W.M., Y.H., and J.X. wrote the manuscript. S.X. revised the manuscript.

## Supplementary Figures

**Supplementary Fig. 1 *β-ACTIN-8xMS2* mRNA localized to the lamellipodia.**

Confocal image of *β-ACTIN-8xMS2* mRNA in live HeLa cells. sfGFP foci are shown. Constructs of *β-ACTIN-8xMS2*, MCP-24xSuntag, and scFv-sfGFP were cotransfected into HeLa cells. Images were taken 12 hr after transfection. The white dashed line demarcates the cell. The white arrow indicates lamellipodia. Scale bar, 5 μm.

**Supplementary Fig. 2 Live imaging of *cdc42* mRNA in *C. elegans* using MS2 based signal amplification with Suntag System.**

Representative images of *cdc42-8xMS2* mRNA in live *C. elegans*. sfGFP fluorescence signals are shown. Left panel: Constructs of *cdc42-8xMS2*, MCP-24xSuntag, and scFv-sfGFP were co-expressed in the epidermis of *C. elegans*. Right panels: where one of the elements was removed as indicated. Scale bar, 10 μm.

**Supplementary Fig. 3 Live imaging of endogenous mRNA in the epidermis of *C. elegans* using the MS2 based signal amplification with Suntag System.**

**a**, Representative images of endogenous *mai-1-8xMS2* mRNA in the epidermis of live *C.elegans* using the strategy described in **Fig. 2a**. Left: *mai-1* without 8xMS2. Right: *mai-1* with 8xMS2. Images were taken before and 15 min after wounding. White dashed circles indicate the wound sites. Scale bar, 10 μm.

**b**, Quantitative RT-PCR showing the expression level of endogenous *mai-1* mRNA in *C. elegans* before and 15 min after wounding. *n* = 3 independent experiments; bars indicate mean ± SEM. Mann-Whitney test, ***p < 0.001.

**Supplementary Fig. 4 Fast activation and spreading of endogenous gene expression in the skin of *C. elegans***

Time-lapse imaging of endogenous *c42d4.3-8xMS2* mRNA in the epidermis of live *C.elegans* before and after wounding. White dashed circles indicate the wound sites. Scale bar, 10 μm.

**Supplementary Fig. 5 Treatment of Actinomycin D blocks the formation of *c42d4.3* mRNA foci**

Time-lapse imaging of endogenous *c42d4.3-8xMS2* mRNA in the epidermis of live *C.elegans* before and after wounding. *C. elegans* were treated with Actinomycin D (30 μM) for 3 hours before wounding. White dashed circles indicate the wound sites. Scale bar, 10 μm.

## Supplementary Videos

**Supplementary Video 1**

Time-lapse imaging of *β-ACTIN-8xMS2* mRNA in live HeLa cells. The sfGFP foci of the *β-ACTIN-3’UTR-8XMS2* mRNA localize to the lamellipodia in HeLa cells. The white dashed line demarcates the cell. The white arrow indicates lamellipodia. Time interval, 2 seconds. Scale bar, 5 μm.

**Supplementary Video 2**

Time-lapse imaging of sfGFP foci of *β-ACTIN-8XMS2* mRNA with a time interval of 1 second in HeLa cells. Scale bar, 5 μm.

**Supplementary Video 3**

Time-lapse imaging showing fusion events of sfGFP foci of *β-ACTIN-8XMS2* mRNA in HeLa cells where small sfGFP spots (white and yellow arrows) fuse into a single more prominent spot. Time interval, 2 seconds. Scale bar, 1 μm.

**Supplementary Video 4**

Time-lapse imaging showing fission events of sfGFP foci of *β-ACTIN-8XMS2* mRNA in HeLa cells where large sfGFP foci split into smaller spots (white and yellow arrows). Time interval, 1 second. Scale bar, 1 μm.

**Supplementary Video 5**

Time-lapse imaging showing transient interactions of an sfGFP spot (yellow arrow) between two spots (white arrows) of *β-ACTIN-8XMS2* mRNA in HeLa cells. Time interval, 2 seconds. Scale bar, 1 μm.

**Supplementary Video 6**

Time-lapse imaging in HeLa cells showing an sfGFP spot of *β-ACTIN-8XMS2* mRNA showing no movement over a 10 second period. Time interval, 1 second. Scale bar, 1 μm.

**Supplementary Video 7**

Time-lapse imaging of endogenous *c42d4.3-8xMS2* mRNA dynamics in the epidermis of *C.elegans* after laser wounding. Time interval, 5 seconds. Scale bar, 10 μm.

**Supplementary Video 8**

Time-lapse imaging showing fusion events of sfGFP foci of endogenous *c42d4.3-8xMS2* mRNA in the epidermis of *C. elegans* where small sfGFP spots (white arrows) fused into a more prominent spot. Time interval, 5 seconds. Scale bar, 5 μm.

## Methods

### Cell lines

The HEK293T/17 cell line was purchased from Procell. The human cervical cancer cell line, HeLa, was a gift from the lab of Christine Mayr (Memorial Sloan Kettering Cancer Center). Cells were maintained at 37°C with 5% CO2 in Dulbecco’s Modified Eagle Medium (DMEM) containing 4,500 mg/L glucose, 10% fetal bovine serum, 100 U/ml penicillin, and 100 mg/ml streptomycin.

### Worm culture

All strains were cultured on the nematode growth medium (NGM) plates with E. coli OP50 at 20-22.5°C, unless otherwise indicated. The N2 Bristol strain was used as the wild-type strain.

### Constructs for mammalian cells

#### MS2 constructs

2xMS2 was designed based on the MS2 sequence as reported by the Singer Lab (Addgene #27118). The sequence of 2xMS2 was as follows: ctgcaggtcgactctagaaaacatgaggatcacccatgtctgcaggtcgactctagaaaacatgaggatcacccatgt. EcoRI-2xMS2-EcoRV was synthesized and inserted into a pcDNA-puro-BFP backbone with EcoRI and EcoRV sites to make pcDNA-puro-BFP-2xMS2. EcoRV-2xMS2-XhoI was synthesized and inserted into a pcDNA-puro-BFP-2xMS2 backbone with EcoRV and XhoI sites to make pcDNA-puro-BFP-4xMS2. XhoI-2xMS2-ApaI was synthesized and inserted into pcDNA-puro-BFP-4xMS2 with XhoI and ApaI sites to make pcDNA-puro-BFP-6xMS2. BamHI-2xMS2-EcoRI was synthesized and inserted into pcDNA-puro-BFP-6xMS2 with BamHI and EcoRI sites to make pcDNA-puro-BFP-8xMS2.

To make pcDNA-puro-BFP-*β-ACTIN*-3’UTR-8xMS2, a *β-ACTIN* coding sequence with a 3’UTR of 373 bp was PCR-amplified from the cDNA of HEK293T/17 cells and cloned into pcDNA-puro-BFP-8xMS2 with KpnI and BamHI sites.

To make pcDNA-puro-BFP-*β-ACTIN*-3’UTR, a *β-ACTIN* coding sequence with a 3’UTR of 373 bp was PCR-amplified from the cDNA of HEK293T/17 cells and cloned into pcDNA-puro-BFP with KpnI and BamHI sites.

#### Suntag and MCP constructs

To make pcDNA-puro-MCP-mCherry-24xSuntag, 24xSuntag (24xGCN4) was PCR-amplified from pcDNA4TO-24xGCN4_v4-kif18b-24xPP7 (Addgene, #74928) and inserted between the BsrGI and ApaI sites of pcDNA-MCP-mCherry.

To make pcDNA-puro-mCherry-24xSuntag, mCherry-24xSuntag was PCR-amplified from pcDNA-puro-MCP-mCherry-24xSuntag. MCP-mCherry was then cut off from the pcDNA-puro-MCP-mCherry backbone by KpnI and XbaI digestion and replaced with mCherry-24xSuntag.

To make pcDNA-puro-scFv-sfGFP, scFV-HAtag-sfGFP-GBI was PCR-amplified from pHR-scFv-GCN4-sfGFP-GB1-dWPRE (Addgene, #60907) and inserted between the NheI and HindIII sites of the pcDNA-puro vector.

To make pcDNA-puro-sfGFP, HAtag-sfGFP-GBI was PCR-amplified from pcDNA-puro-scFv-sfGFP and inserted between the NheI and HindIII sites of the pcDNA-puro vector.

#### Constructs for *C. elegans* and transgenic worms

The elements of the MS2-Suntag system were cloned by PCR from the plasmids described above and were inserted into vectors for expression in *C. elegans*.

Different combinations of plasmids were used for generating extrachromosomal array transgenic worms. Briefly, 50 ng/μl plasmids of P*semo-1*-MCP-24×Suntag, 10 ng/μl plasmids of P*col-19*-antibody-sfGFP and 10 ng/μl co-injection marker *Pttx-3-RFP* were injected into N2 and knock-in animals. The extrachromosomal strains were as follows: SHX3314: *Pcol-19*-mKate2::CDC-42-8×MS2, P*semo-1*-MCP-24×Suntag, *Pcol-19-antibody-sfGFP(zjuEx2144);* SHX3749: P*col-19*-mKate2::CDC-42-8×MS2; P*semo-1*-MCP; P*col-19*-antibody-*sfGFP(zjuEx2031);* SHX3751: P*col-19*-mKate2::CDC-42-8×MS2; P*semo-1*-MCP-24×Suntag; *Pcol-19-sfGFP(zjuEx2033);* SHX3755: P*col-19*-mKate2::CDC-42-8×MS2; P*semo-1*-24×Suntag; P*col-19*-antibody-sfGFP(*zjuEx2037)*; SHX3757: P*col-19*-mKate2::CDC-42; P*semo-1*-MCP-24×Suntag; P*col-19*-antibody-sfGFP (*zjuEx2144);* SHX3908: *c42d4.3-8×MS2(syb5468);* P*semo-1*-MCP-24×Suntag; P*col-19*-antibody-sfGFP(*zjuEx2144);* SHX3909: *mai-1-8×MS2(syb5458);* P*semo-1*-MCP-24×Suntag; and P*col-19*-antibody-sfGFP(*zjuEx2144)*.

#### Transfections

Lipofectamine 2000 (Invitrogen) was used for all transfections.

#### Live cell imaging

HeLa cells were plated on 35 mm glass-bottom dishes (NEST, 801001) at a density of 0.13×10^6^. The indicated constructs were transfected into HeLa cells. 12-24 hours after transfection, cells were imaged using a spinning-disk confocal microscope with a 60×objective (Nikon T2 Microscope; Apo 60x oil; 1.4 NA) using the SoRa mode. Exposure time was 500 ms. For time-lapse imaging, the time interval was set to 1s or 2s. Images were analyzed with FIJI (ImageJ).

#### CRISPR-Cas9 mediated gene knock-in in *C. elegans*

*C42d4.3-8×MS2(syb5468)* and *mai-1-8×MS2(syb5458)* knock-in animals were generated using the CRISPR-Cas9 system. Briefly, two repair templates were cloned using the Gibson assembly technique within a pDD282 plasmid. The MS2 sequence was designed after the termination codon of two genes in the repair template. sgRNA, repair templates, and *Peft-3-Cas9-NLS-pU6-dpy-10* sgRNA, as well as *Pmyo-2-cherry* as co-injection markers, were injected into N2 worms. Knock-in animals were confirmed by PCR genotyping and sequencing. Roller or dumpy animals were heat-shocked or outcrossed to remove markers. The sequences of the sgRNAs used in this study were as follows: *c42d4.3* sg1 (CCAAAACTTGCTTGCCAGAACTT); *c42d4.3* sg2 (CCAGAACTTTCGGACAATAATTG); *mai-1* sg1 (ACAACATCAGCAACGACTGAAGG); *mai-1* sg2 (CGACTGAAGGAAATCGAGAAAGG). The precise sequence knock-in is described as follows: gggaggtgatagcattgcttggatccctgcaggtcgactctagaaaacatgaggatcacccatgtctgcaggtcgac tctagaaaacatgaggatcacccatgtgaattcctgcaggtcgactctagaaaacatgaggatcacccatgtctgcaggtcgactcta gaaaacatgaggatcacccatgtgatatcctgcaggtcgctctagaaaacatgaggatcacccatgtctgcaggtcgactctagaa aacatgaggatcacccatgtctcgagctgcaggtcgactctagaaaacatgaggatcacccatgtctgcaggtcgactctgaaaac atgaggatcacccatgtgggcccgtttaaacccgctga.

#### Drug treatment

The Actinomycin D stock solution was dissolved in DSMO and diluted with M9 to a working concentration of 30 μM. Young adult stage worms were incubated in 100 μl Actinomycin D (APExBIO; Catalog No. A4448) solution (containing *E. coli* OP50) using a 1.5 ml microcentrifuge tube at 20 °C for 3 hours. The worms were then transferred to fresh NGM plates to dry before wounding and imaging.

#### Wounding assay

A Micropoint UV laser was used to wound the epidermis of young adult stage worms. Briefly, worms at the young adult stage were mounted to 4% agarose gel on a slide, narcotized with 12 μM Levamisole, and wounded using a Micropoint UV laser.

#### Live Imaging in *C. elegans*

Worms were imaged on a spinning disk confocal microscope (Andor 100x, NA 1.46 objective). Z-stack and time-lapse were set using Andor IQ software to capture the images.

#### Quantification of the size and number of sfGFP foci

The size and number of sfGFP were quantified using Fiji software (https://imagej.net/imagej-wiki-static/Fiji). The background intensity was set as the threshold, and the size and number of foci were calculated using the Analyze Particles command. The value of puncta size and number were statistically analyzed using GraphPad. For puncta size and number, we chose 600 × 400-pixels around the wound site for analysis. The size and number changes were quantified using mean with Standard Error of the Mean (SEM).

#### qRT-PCR

Total RNAs were extracted from 100 young adult worms with TRIzol reagent (Invitrogen, Carlsbad, CA, USA), quantitated by spectrophotometry using a NanoDrop (Thermo, USA), and reverse transcribed using HiScript®IIIReverseTranscriptase (Vazyme, China). qRT-PCR was performed with *rbd-1* as the house-keeping gene using the SYBR Green Supermix (Vazyme). The following primers were used in this study.

*rbd-1*, forward (fwd) CACGGAACAGCAACTACGGA, reverse (rev) CGGCTTGTTTGCATCACCAA;

*c42d4.3*, fwd GCCAGACTCTTGCCTCTCAA, rev CACGCGGTGTGATCTTTTCC;

*mai-1*, fwd CGGCTCAATCCGTGAAGC, rev TGTTGGCTTTGCGTCATATC-3

#### Statistical analysis

Statistical analyses were performed using GraphPad Prism 7 (La Jolla, CA). A non-parametric Mann-Whitney test was used for two comparisons. Ns donates no significant difference; *** indicates *P* < 0.001. Unless elsewhere stated, bars represent means ± SEM.

## Reference

1. Buxbaum, A.R., Haimovich, G. & Singer, R.H. In the right place at the right time: visualizing and understanding mRNA localization. Nature Reviews Molecular Cell Biology 16, 95–109 (2015).

2. Braselmann, E., Rathbun, C., Richards, E.M. & Palmer, A.E. Illuminating RNA Biology: Tools for Imaging RNA in Live Mammalian Cells. Cell chemical biology 27, 891–903 (2020).

3. Le, P., Ahmed, N. & Yeo, G.W. Illuminating RNA biology through imaging. Nature Cell Biology (2022).

4. Bertrand, E. et al. Localization of ASH1 mRNA particles in living yeast. Mol Cell 2, 437–445 (1998).

5. Nelles, David A. et al. Programmable RNA Tracking in Live Cells with CRISPR/Cas9. Cell 165, 488–496 (2016).

6. Yang, L.-Z. et al. Dynamic Imaging of RNA in Living Cells by CRISPR-Cas13 Systems. Molecular Cell 76, 981–997.e987 (2019).

7. Chen, X. et al. Visualizing RNA dynamics in live cells with bright and stable fluorescent RNAs. Nature Biotechnology 37, 1287–1293 (2019).

8. Paige, J.S., Wu, K.Y. & Jaffrey, S.R. RNA mimics of green fluorescent protein. Science (New York, N.Y.) 333, 642–646 (2011).

9. Sunbul, M. et al. Super-resolution RNA imaging using a rhodamine-binding aptamer with fast exchange kinetics. Nat Biotechnol 39, 686–690 (2021).

10. Halstead, J.M. et al. Translation. An RNA biosensor for imaging the first round of translation from single cells to living animals. Science (New York, N.Y.) 347, 1367–1671 (2015).

11. Levo, M. et al. Transcriptional coupling of distant regulatory genes in living embryos. Nature 605, 754–760 (2022).

12. Dufourt, J. et al. Imaging translation dynamics in live embryos reveals spatial heterogeneities. Science (New York, N.Y.) 372, 840–844 (2021).

13. Park, H.Y. et al. Visualization of dynamics of single endogenous mRNA labeled in live mouse. Science (New York, N.Y.) 343, 422–424 (2014).

14. Das, S., Moon, H.C., Singer, R.H. & Park, H.Y. A transgenic mouse for imaging activity-dependent dynamics of endogenous Arc mRNA in live neurons. Science advances 4, eaar3448 (2018).

15. Zimyanin, V.L. et al. In Vivo Imaging of oskar mRNA Transport Reveals the Mechanism of Posterior Localization. Cell 134, 843–853 (2008).

16. Tanenbaum, M.E., Gilbert, L.A., Qi, L.S., Weissman, J.S. & Vale, R.D. A protein-tagging system for signal amplification in gene expression and fluorescence imaging. Cell 159, 635–646 (2014).

17. Katz, Z.B. et al. β-Actin mRNA compartmentalization enhances focal adhesion stability and directs cell migration. Genes & development 26, 1885–1890 (2012).

18. Wang, J. et al. Efficient targeted insertion of large DNA fragments without DNA donors. Nature methods 19, 331–340 (2022).

19. Xu, S. & Chisholm, A.D. Methods for skin wounding and assays for wound responses in C. elegans. Journal of visualized experiments : JoVE (2014).

20. Guo, Y. & Lee, R.E.C. Long-term imaging of individual mRNA molecules in living cells. Cell Reports Methods.

