## Supplementary figures for "Enhanced Single RNA Imaging Reveals Dynamic Gene Expression in Live Animals"

Hu et al. Supplementary Fig. 1

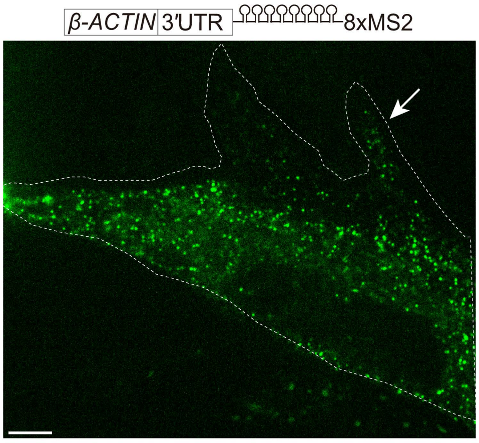


**Supplementary Fig. 1 *β-ACTIN-8xMS2* mRNA localized to the lamellipodia.**
Confocal image of *β-ACTIN-8xMS2* mRNA in live HeLa cells. sfGFP foci are shown. Constructs of *β-ACTIN-8xMS2*, MCP-24xSuntag, and scFv-sfGFP were cotransfected into HeLa cells. Images were taken 12 hr after transfection. The white dashed line demarcates the cell. The white arrow indicates lamellipodia. Scale bar, 5 μm.

Hu et al. Supplementary Fig. 2

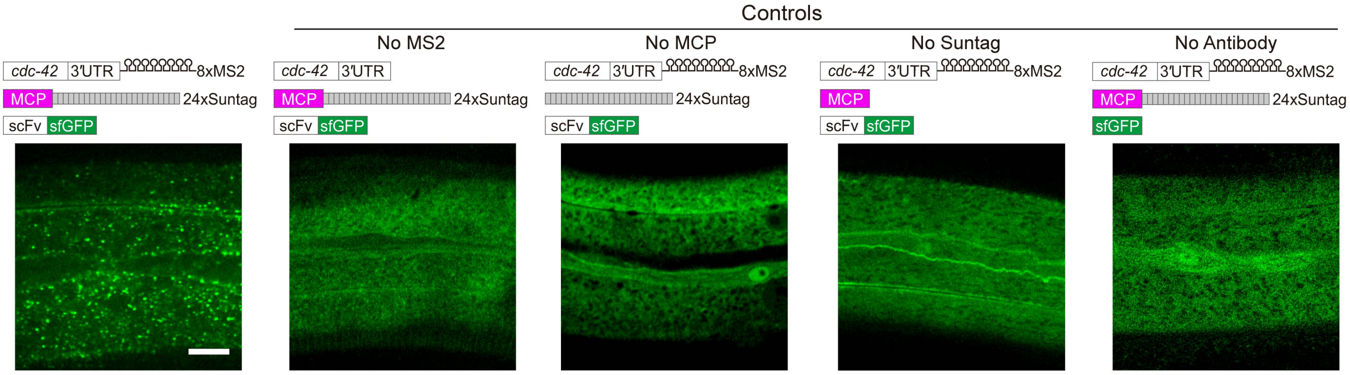


**Supplementary Fig. 2 Live imaging of *cdc42* mRNA in *C. elegans* using the MS2 based signal amplification with Suntag system.** Representative images of *cdc42-8xMS2* mRNA in live *C. elegans*. sfGFP fluorescence signals are shown. Left panel: Constructs of *cdc42-8xMS2*, MCP-24xSuntag, and scFv-sfGFP were co-expressed in the epidermis of *C. elegans*. Right panels: where one of the elements was removed as indicated. Scale bar, 10 μm.

Hu et al. Supplementary Fig. 3

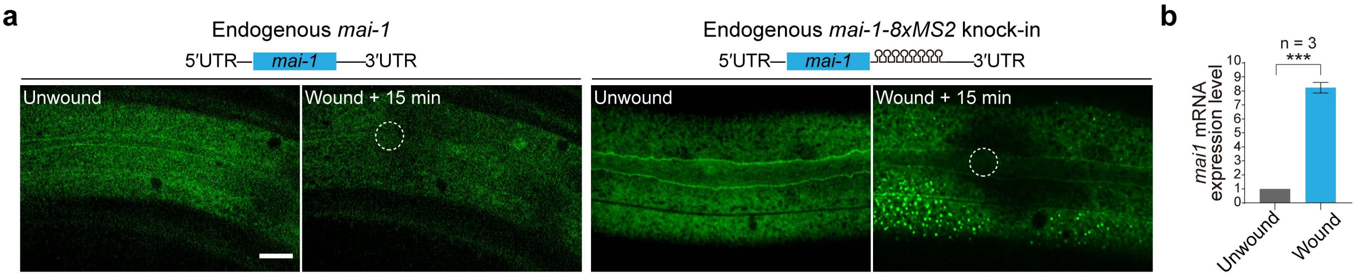


**Supplementary Fig. 3 Live imaging of endogenous mRNA in the epidermis of *C. elegans* using the MS2 based signal amplification with Suntag system.**

**a**, Representative images of endogenous *mai-1-8xMS2* mRNA in the epidermis of live *C.elegans* using the strategy described in **Fig. 2a**. Left: *mai-1* without 8xMS2. Right: *mai-1* with 8xMS2. Images were taken before and 15 min after wounding. White dashed circles indicate the wound sites. Scale bar, 10 μm.

**b**, Quantitative RT-PCR showing the expression level of endogenous *mai-1* mRNA in *C. elegans* before and 15 min after wounding. n = 3 independent experiments; bars indicate mean ± SEM. Mann-Whitney test, ***p < 0.001.

Hu et al. Supplementary Fig. 4

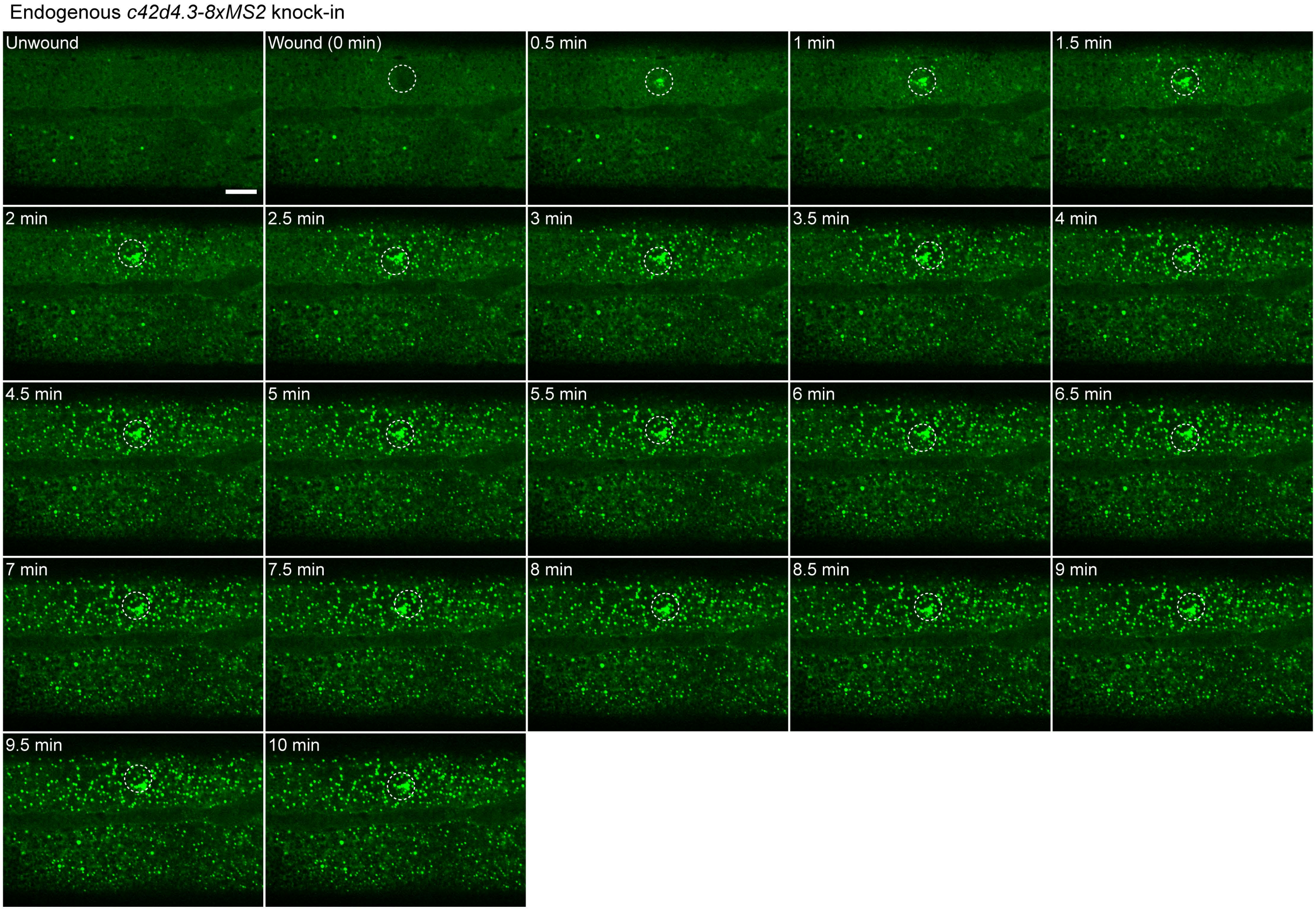


**Supplementary Fig. 4 Fast activation and spreading of endogenous gene expression in the skin of *C. elegans*.**
Time-lapse imaging of endogenous *c42d4.3-8xMS2* mRNA in the epidermis of live *C.elegans* before and after wounding. White dashed circles indicate the wound sites. Scale bar, 10 μm.

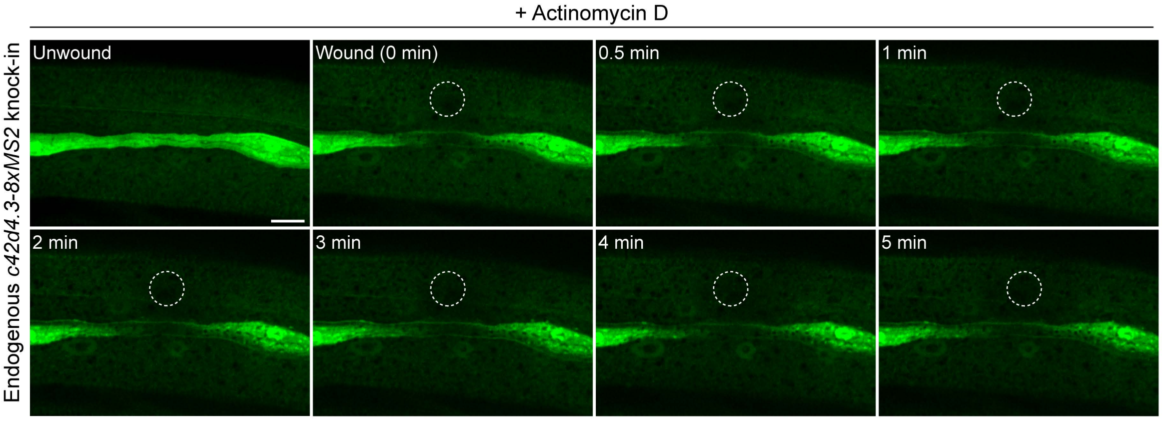


**Supplementary Fig. 5 Treatment of Actinomycin D blocks the formation of *c42d4.3* mRNA foci.**
Time-lapse imaging of endogenous *c42d4.3-8xMS2* mRNA in the epidermis of live *C.elegans* before and after wounding. *C. elegans* were treated with Actinomycin D (30 μM) for 3 hours before wounding. White dashed circles indicate the wound sites. Scale bar, 10 μm.
